# Reversal of molecular pathology by RNA-targeting Cas9 in a myotonic dystrophy mouse model

**DOI:** 10.1101/184408

**Authors:** Ranjan Batra, David A. Nelles, Florian Krach, James D. Thomas, Lukasz Snjader, Steven M. Blue, Stefan Aigner, Maurice S. Swanson, Gene W. Yeo

## Abstract

The dominantly inherited, multi-systemic disease myotonic dystrophy type I (DM1) is caused by triplet repeat CTG expansions in the *DMPK* gene and is the most common form of adult-onset muscular dystrophy. Elimination of the toxic, repetitive CUG RNA constitutes a therapeutic for this disease. We report an RNA-targeting Cas9 (RCas9) system that supports efficient reversal of DM1 phenotypes via delivery to adult poly(CUG) DM1 mouse muscle using adeno-associated virus (AAV). We observe elimination of CUG RNA, restoration of CUG foci-associated Mbnl1 protein to wild-type subcellular localization, correction of DM1-type alternative splicing patterns in candidate genes including the voltage-gated chloride channel 1 (*Clcn1)* responsible for characteristic myotonia, recovery of Clcn1 staining, and reduction in centralized myonuclei. Our results establish RCas9 as a potential long-term *in vivo* therapeutic for DM1.

**One Sentence Summary:** A repurposed CRISPR system termed RNA-targeting Cas9 reverses the molecular pathology associated with the most common type of adult onset muscular dystrophy in adult mouse muscle.

## Main text

Myotonic dystrophy type 1 is an autosomal inherited disorder characterized by CTG repeat expansions in the *DMPK* gene. The repetitive RNAs produced by this locus form nuclear RNA foci (1) that sequester RNA binding proteins such as MBNL1 (2, 3) and divert them from their homeostatic RNA processing activities (4, 5). Resulting loss of MBNL1 function is linked to hundreds of splicing defects that cause myotonia and progressive muscle degeneration (4, 6, 7). Since *DMPK* is widely expressed, DM1 pathology is multi-systemic leading to subcapsular cataracts, cognitive defects, and cardiac conduction defects that contribute to DM1-associated mortality (8). Currently available treatments for DM1 do not address the underlying etiology of this disease that affect more than 30,000 patients in the US alone and a recent clinical failure of a promising antisense oligonucleotide (ASO) targeting CUG repeat RNA (clinicaltrials.gov identifier: NCT02312011) highlights the need for new therapeutic modalities.

Technologies that engage the DM1-linked repeat expansion on both the DNA and RNA levels have been investigated. Although excision of these large tracts in DNA followed by repair in their absence is possible (9), this genome engineering approach has not yet been realized with high efficiency. ASOs provide means to engage pathogenic RNAs directly but must be continuously re-administered for life and have experienced poor biodistribution in muscle in the clinic. RNA interference (RNAi) can be encoded in adeno-associated viral vectors to support long-term, targeted delivery to human tissue (10) but typically does not engage repetitive RNAs efficiently.

By repurposing the CRISPR/Cas9 genome engineering system, we have recently demonstrated the ability of Cas9 to efficiently engage RNA (11) and eliminate microsatellite repeat expansion RNAs [Batra et al, 2017] in human cells. Utilizing a nuclease-null Cas9 (dCas9) fused to an RNA endonuclease, this RNA-specific CRISPR system features no inherent risk of off-target genome editing (12). In this study, we set out to investigate whether AAV-packaged RNA-targeting Cas9 (RCas9) supports elimination of DM1-linked repetitive CUG^exp^ RNA *in vivo* after delivery to adult mouse muscle.

Mouse models that express CUG repeat RNA in various tissues (poly(CUG)) recapitulate DM1 phenotypes (13) and have been used extensively to evaluate efficacy of various therapeutic approaches such as ASOs (14–16) and Mbnl1 gene therapy (4). Mbnl1 knockout models feature similar phenotypes in muscle (17) as poly(CUG) binds and sequesters Mbnl1 that lead to disruption of its physiological alternative splicing and polyadenylation activities. This disruption of Mbnl1-mediated RNA processing is directly linked to hallmark phenotypes including myotonia. We packaged the RNA-targeting Cas9 system composed of dCas9 fused to the PIN RNA endonuclease in AAV-9 and constructed a second vector carrying a U6 promoter-driven single guide RNA (sgRNA) (18) with CMV-driven GFP (Figure 1A). rAAV-9 containing PIN-dCas9 was combined with either rAAV9 carrying CUG-targeting (RCas9-CUG) or non-targeting sgRNA (RCas9-NT) and injected (2.5x 10^10^ vg each) into contralateral tibialis anterior (TA) muscles of 8-week old HSA_LR_ poly(CUG) mice (N=3) that express 250 CTG repeats downstream of the human skeletal actin (HSA) promoter in muscle (Figure 1B) (13). TA was harvested 3 weeks post-injection with no obvious signs of inflammation (data not shown). We observed efficient elimination of the characteristic CUG repeat RNA foci by RNA dot blot using CAG^10^ probes (Figure S1A) and RNA fluorescence in situ hybridization (FISH) (Figure 1C-D, N=3 mice with contralaterally-injected CUG-targeting and non-targeting RCas9 systems) in the muscle injected with the CUG-targeting RCas9 system. This result is consistent with previous results in DM1 patient myoblasts that indicate similar sequence-specific elimination of CUG repeat RNA [Batra et al 2017]. Muscle treated with the non-targeting RCas9 system and contralateral saline control featured similar levels of RNA foci.

**Fig. 1.**
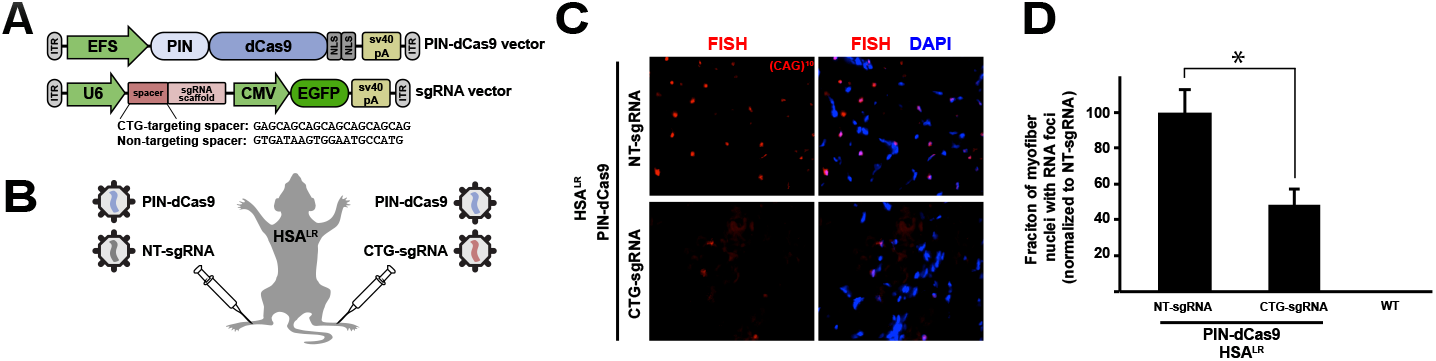
Elimination of CUG repeat RNA in a mouse model of myotonic dystrophy type 1. (A) Schematic of the structure of the PIN-dCas9 and sgRNA AAV vectors. (B) Schematic description of contralateral injections of CUG-targeting and non-targeting RCas9 systems into the tibialis anterior (TA) muscles of HSA_LR_ mice. (C-D) RNA FISH for CUG repeat RNA in tibialis anterior muscle subjected to the CUG-targeting and nontargeting RCas9 system. Error bars indicate standard deviation (N=3 mice with contralateral targeting and non-targeting RCas9 systems. Measurements were conducted on 3 serial muscle sections each)

Next, we set out to evaluate the consequence of this efficient and specific elimination of CUG repeat RNA foci on downstream DM1 phenotypes. The MBNL family of proteins comprises RNA binding proteins that are responsible for developmental regulation of hundreds of RNA processing events in various tissues (5, 19–21). Loss of MBNL function through sequestration by CUG^exp^ leads to mis-splicing of many of its target RNAs including inclusion of exon 7a (contains a premature termination codon) in chloride voltage-gated channel 1 (*CLCN1)* which leads to non-sense mediated decay. Resulting reduction in CLCN1 protein levels causes DM1’s namesake myotonia. To evaluate the potential of RCas9 to reverse this DM1-linked spliceopathy, we first assessed the localization of MBNL1 protein in mouse muscle tissue and observed that tissue treated with the CUG-targeting RCas9 system exhibited diffuse nuclear MBNL1 localization similar to wild-type mice (Figure 2A). To assess whether known MBNL1-mediated splicing defects were ameliorated, we conducted RT-PCR with primers flanking alternatively spliced exons in *Clcn1* (Exon 7a), *Atp2a1* (Exon 22), *Tnnt3* (Exon F), and *Ldb3* (also called *Zasp* or *Cypher,* Exon 11; Figure 2B; N=3 mice with contralaterally-injected CUG-targeting and non-targeting RCas9 systems). CUG-targeting RCas9 promoted reversal of splicing defects to resemble wild type patterns while contralateral muscles injected with a non-targeting RCas9 system resembled known

**Fig. 2.**
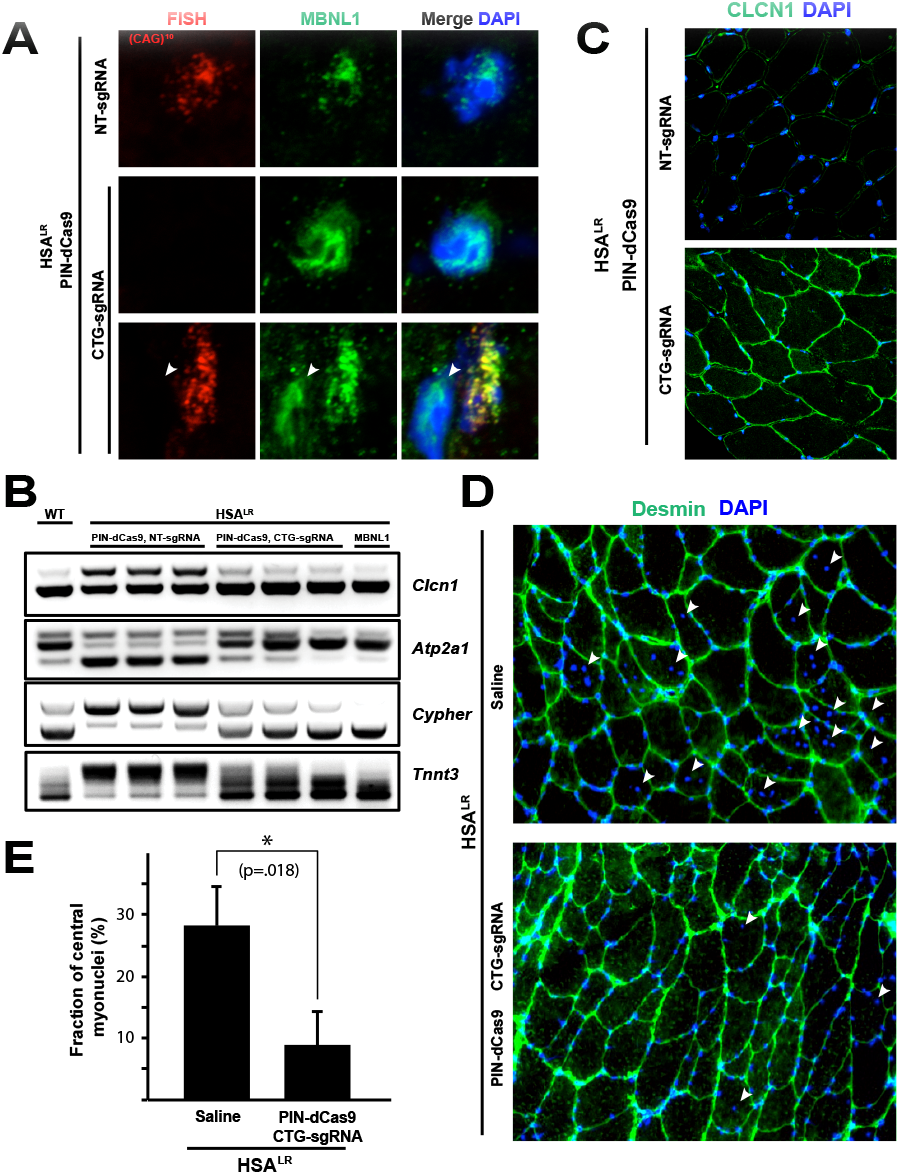
RNA-targeting Cas9 reverses myotonic dystrophy type I pathology in adult mouse muscle. (A) Mbnl1 protein and CUG repeat RNA foci localization are visualized by immunofluorescence and RNA FISH, respectively. In the bottom panel, a cell lacking CUG repeat RNA is highlighted for comparison with an adjacent CUG repeat-harboring cell. (B) Splicing of alternative exons regulated by Mbnl1 (*Clcn1, Atp2a1, Ldb3, and Tnnt3*) was assessed by RT-PCR. Rescue of disease-associated splicing is observed by the RCas9 system (PIN-dCas9, CTG-sgRNA) to resemble wildtype (WT) patterns. Total RNA was extracted from N=3 mice with contralateral injections of targeting and nontargeting RCas9 systems. (C) Levels of *Clcn1* protein are assessed by immunofluorescence in transverse sections of tibialis anterior mouse muscle in the presence of the CUG RNA-targeting (CTG-sgRNA) and non-targeting (NT-sgRNA) RCas9 systems. (D) Central nucleation in muscle fibers was assessed in transverse sections and quantified (E). Error bars indicate standard deviation (N=3 mice with contralateral injection of targeting and non-targeting RCas9 systems. Measurements were conducted on 3 serial muscle sections each)

DM1 splicing patterns (4). These results indicate that RCas9 supports reversal of hallmark molecular defects associated with DM1 in adult mouse muscle.

Mis-splicing of *CLCN1* via inclusion of exon 7a causes loss of *CLCN1* protein via inclusion of a premature termination codon or generation of a dominant-negative truncated protein (22, 23). HSA^LR^ mice show reduced Clcn1 staining in muscle (13) which correlates with myotonia. We observed that CUG-targeting RCas9 rescues this loss of Clcn1 protein by immunofluorescence (Figure 2C) indicating rescue of important gene products relevant to DM1 pathology. Muscle damage and resulting turnover of myofibers in HSA_LR_ causes increased fraction of central nucleation (4). This hallmark of ongoing muscle regeneration is reduced by the RCas9 system (Figure 2D-E, N=3 mice with contralaterally-injected CUG-targeting and non-targeting RCas9 systems), indicating that muscle turnover is reduced. Motivated by these efficient reversals of hallmark DM1 molecular phenotypes, we next set out to assess transcriptome-wide rescue of alternative splicing and off-target effects of the RCas9 system.

Previous work involving RCas9 targeting poly(CUG) in patient myoblasts indicated that RCas9 corrected ~93% of all mis-splicing events in DM1 myotubes [Batra et al, 2017]. We conducted an analogous measurement in mouse muscle by RNA-seq. Consistent with the RCas9 system in human DM1 myotubes, the RCas9 system in mouse skeletal muscle promoted at least partial reversal of 86% of all DM1-related mis-splicing events (61% full reversal and 25% partial reversal, Figure 3A-B, Supplementary Table 1, N=3 mice with contralaterally-injected CUG-targeting and non-targeting RCas9 systems), resulting in the hierarchical clustering of transcriptome-wide alternative splicing changes with Mbnl1-treated muscle (Figure 3A). Genome browser track of *Clasp1* shows reversal of exon 20 mis-splicing both in the presence of RCas9 and Mbnl1 overexpression (Figure 3C). Furthermore, muscle-specific gene expression was specifically upregulated (Figure 3D, Supplementary Table 2E) in both RCas9- and Mbnl1-treated TA muscle. These results indicate that RCas9 supports highly efficient reversal of the major molecular defects associated with DM1 and induces expression of muscle markers, indicating reversal of dysfunctional muscle differentiation associated with the disease (24).

**Fig. 3.**
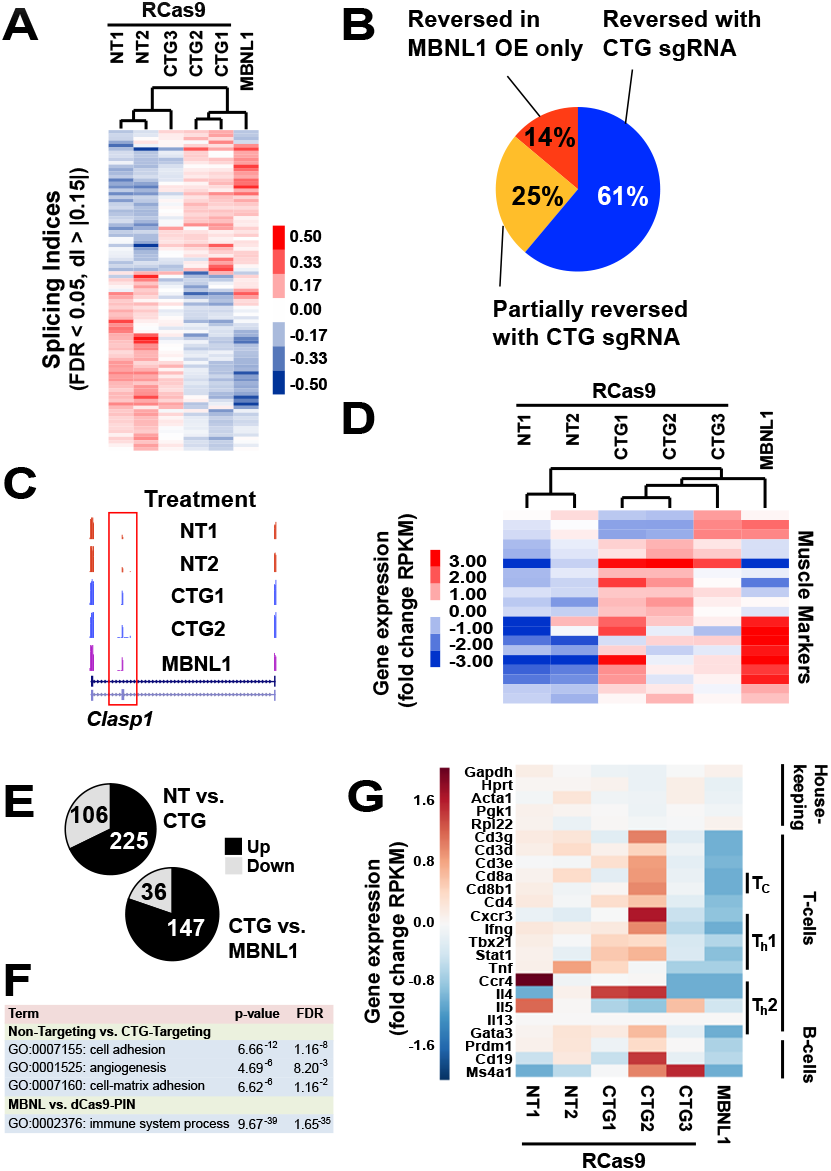
Broad reversal of disease-associated alternative splicing patterns and enhanced expression of mature muscle markers in the presence of the RCas9 system. (A) Hierarchical clustering of splicing indices of genes most altered in HSA_LR_ TA muscle compared to a healthy control. Tissues were treated with either CUG repeat-targeting RCas9 system (“CTG1”, “CTG2”, or “CTG3”), non-targeting RCas9 system (“NT1” or “NT2”), or Mbnl1 protein (“MBNL1”). Each numbered sample represents a mouse (contralateral targeting and non-targeting systems, N=3 mice). (B) Summary of DM1-related alternative splicing reversal describing fraction of genes reversed by the RCas9 system compared to all genes regulated by MBNL1 overexpression. (C) UCSC genome-browser tracks of *Clasp1* showing reversal of DM1-type alternative splicing of exon 20 by both CUG-targeting RCas9 and Mbnl1 overexpression. (D) Expression levels of mature muscle markers in tissues treated with the RCas9 system of Mbnl1 protein. The CUG-targeting RCas9 system and Mbnl1 cluster together and display similar increases in muscle marker expression compared to the non-targeting system. Sample labeling is identical to that in (A). (E) Pie charts showing gene expression levels among the CUG-targeting and non-targeting RCas9 systems or the CUG-targeting RCas9 system and MBNL1 reveal groups of concomitantly increasing (“up”) and decreasing (“down”) genes among these pairs. (F) Gene ontology (GO) analysis comparing CUG-targeting and non-targeting RCas9 systems or the CUG-targeting RCas9 system and MBNL1 reveal distinct groups of gene expression differences (G) Expression of genes linked to adaptive immunity comparing in muscle tissue treated with the CUG-targeting RCas9 system, non-targeting system, or MBNL1 overexpression. Sample labeling is identical to that in (A).

RNA-targeting therapeutics that rely on nucleic acid base-pairing run the inherent risk of off-target alterations to the transcriptome. We have previously shown that elimination of CUG repeat RNAs in human patient myoblasts incurs minimal off-target effects on gene expression and is specific to large repetitive RNAs while leaving short, non-pathogenic repeats unaffected. To assess the off-target effects of RCas9 *in vivo,* we evaluated transcriptome-wide changes and found hundreds of genes featuring expression level changes (Figure 3E, Supplementary Table 2). Strikingly, gene ontology (GO) analysis (Supplementary Table 3) using DAVID (25) showed that the top GO categories representing gene expression changes between targeting and non-targeting RCas9 system were “cell adhesion”, “angiogenesis”, and “cell-matrix adhesion” (Figure 3F). The lack of off-target effects observed *in vitro* and known defects related to cell adhesion, extracellular matrix adhesion (7), and myogenesis (26) in DM1 indicate that these changes are likely due to reversal of DM1-associated pathology by the RCas9 system. We further assessed potential off-target effects by evaluating expression of *Dmpk* which contains 5-20 CTG repeats in HSA_LR_ mice. We observed no differences in *Dmpk* RNA levels (Figure S1B) which is consistent with *in vitro* studies indicating the specificity of RCas9 for long poly(CUG) tracts [Batra et al 2017].

Immune response to Cas9 involving AAV-mediated, long-term expression in tissue is a major consideration for translation of any CRISPR-based therapeutic to the clinic. Recent efforts involving delivery of Cas9 to adult muscle have assessed immune response to varying degrees (27, 28) and detailed assessments have revealed adaptive immune response to Cas9 (29). We assessed gene expression changes between RCas9- and Mbnl-treated mice to assess the immune response to Cas9 protein and observed enrichment in adaptive immune response-linked genes (Figure 3G) and no striking alterations in gene expression linked to innate immune response (Figure S2). Consistent with previous studies (29), we do not observe concomitant changes in muscle morphology and pathology (Figure 2E) in terms of myofiber integrity and myofiber central nucleation. This result indicates that either immune response to the RCas9 system is insufficient to promote significant tissue damage or that some degree of damage is obscured by the improvement in muscle health afforded by the RCas9 system. While the therapeutic effect of RCas9 dominates our observations, future studies should include other animal models that more closely recapitulate the human adaptive immune response.

Microsatellite repeat expansions cause a host of human diseases (30, 31) that frequently arise from toxic gains-of-function at the levels of repetitive RNA and/or aberrant protein. This effort builds upon our recent efforts describing the ability of RCas9 to target and eliminate repetitive RNAs including the CUG repeat linked to DM1. This AAV-encodable system leverages the long-term generation of therapeutic materials that is possible with AAV in contrast to other RNA-engaging therapeutics such as ASOs. While we observe no muscle damage due to the presence of the RCas9 system, a long-term study of efficacy and immune response to the RCas9 system is required. Future studies could also explore the use of muscle-specific promoters or regulatory elements that restrict expression of the RCas9 system to muscle cells. Overall, this effort demonstrates the first prospect of RCas9 as a therapeutic for DM1 and other microsatellite repeat expansion diseases.

## Acknowledgements

Data described in this work are available in the GEO respository (GSEXXXXX). Salary support for G.W.Y. was partially supported by grants from the NIH (HG004659 and NS075449). This work was funded by a start-up grant from the Stem Cell Program at UCSD (G.W.Y). Ranjan Batra is an MDF Postdoctoral Fellow.

## References and Notes

1. J. R. O'Rourke, M. S. Swanson, Mechanisms of RNA-mediated disease. J Biol Chem 284, 7419–7423 (2009).

2. J. W. Miller et al., Recruitment of human muscleblind proteins to (CUG)(n) expansions associated with myotonic dystrophy. The EMBO journal 19, 4439-4448 (2000).

3. J. Shin, K. Charizanis, M. S. Swanson, Pathogenic RNAs in microsatellite expansion disease. Neuroscience letters 466, 99–102 (2009).

4. R. N. Kanadia et al., Reversal of RNA missplicing and myotonia after muscleblind overexpression in a mouse poly(CUG) model for myotonic dystrophy. Proc Natl Acad Sci U S A 103, 11748–11753 (2006).

5. R. Batra et al., Loss of MBNL leads to disruption of developmentally regulated alternative polyadenylation in RNA-mediated disease. Mol Cell 56, 311–322 (2014).

6. T. M. Wheeler, J. D. Lueck, M. S. Swanson, R. T. Dirksen, C. A. Thornton, Correction of ClC-1 splicing eliminates chloride channelopathy and myotonia in mouse models of myotonic dystrophy. The Journal of clinical investigation 117, 3952–3957 (2007).

7. H. Du et al., Aberrant alternative splicing and extracellular matrix gene expression in mouse models of myotonic dystrophy. Nat Struct Mol Biol 17, 187-193 (2010).

8. C. A. Thornton, Myotonic dystrophy. Neurol Clin 32, 705–719, viii (2014).

9. E. L. van Agtmaal et al., CRISPR/Cas9-Induced (CTGCAG)n Repeat Instability in the Myotonic Dystrophy Type 1 Locus: Implications for Therapeutic Genome Editing. Mol Ther 25, 24–43 (2017).

10. L. Naldini, Gene therapy returns to centre stage. Nature 526, 351–360 (2015).

11. D. A. Nelles et al., Programmable RNA Tracking in Live Cells with CRISPR/Cas9. Cell 165, 488–496 (2016).

12. K. A. Schaefer et al., Unexpected mutations after CRISPR-Cas9 editing in vivo. Nat Methods 14, 547–548 (2017).

13. A. Mankodi et al., Myotonic dystrophy in transgenic mice expressing an expanded CUG repeat. Science 289, 1769–1773 (2000).

14. T. M. Wheeler et al., Reversal of RNA dominance by displacement of protein sequestered on triplet repeat RNA. Science 325, 336–339 (2009).

15. J. E. Lee, C. F. Bennett, T. A. Cooper, RNase H-mediated degradation of toxic RNA in myotonic dystrophy type 1. Proc Natl Acad Sci U S A 109, 4221–4226 (2012).

16. T. M. Wheeler et al., Targeting nuclear RNA for in vivo correction of myotonic dystrophy. Nature 488, 111–115 (2012).

17. R. N. Kanadia et al., A muscleblind knockout model for myotonic dystrophy. Science 302, 1978–1980 (2003).

18. B. Chen et al., Dynamic imaging of genomic loci in living human cells by an optimized CRISPR/Cas system. Cell 155, 1479–1491 (2013).

19. X. Lin et al., Failure of MBNL1-dependent post-natal splicing transitions in myotonic dystrophy. Human molecular genetics 15, 2087–2097 (2006).

20. A. Kalsotra et al., A postnatal switch of CELF and MBNL proteins reprograms alternative splicing in the developing heart. Proc Natl Acad Sci U S A 105, 20333–20338 (2008).

21. K. Charizanis et al., Muscleblind-like 2-mediated alternative splicing in the developing brain and dysregulation in myotonic dystrophy. Neuron 75, 437–450 (2012).

22. J. Berg, H. Jiang, C. A. Thornton, S. C. Cannon, Truncated ClC-1 mRNA in myotonic dystrophy exerts a dominant-negative effect on the Cl current. Neurology 63, 2371–2375 (2004).

23. A. Mankodi et al., Expanded CUG repeats trigger aberrant splicing of ClC-1 chloride channel pre-mRNA and hyperexcitability of skeletal muscle in myotonic dystrophy. Mol Cell 10, 35–44 (2002).

24. G. Meola, R. Cardani, Myotonic dystrophies: An update on clinical aspects, genetic, pathology, and molecular pathomechanisms. Biochim Biophys Acta 1852, 594–606 (2015).

25. W. Huang da, B. T. Sherman, R. A. Lempicki, Systematic and integrative analysis of large gene lists using DAVID bioinformatics resources. Nat Protoc 4, 44–57

26. J. D. Amack, M. S. Mahadevan, Myogenic defects in myotonic dystrophy. Dev Biol 265, 294–301 (2004).

27. D. U. Kemaladewi et al., Correction of a splicing defect in a mouse model of congenital muscular dystrophy type 1A using a homology-directed-repair-independent mechanism. Nature medicine, (2017).

28. C. Long et al., Postnatal genome editing partially restores dystrophin expression in a mouse model of muscular dystrophy. Science 351, 400–403 (2016).

29. W. L. Chew et al., A multifunctional AAV-CRISPR-Cas9 and its host response. Nat Methods 13, 868–874 (2016).

30. L. P. Ranum, T. A. Cooper, RNA-mediated neuromuscular disorders. Annual review of neuroscience 29, 259–277 (2006).

31. R. Batra, K. Charizanis, M. S. Swanson, Partners in crime: bidirectional transcription in unstable microsatellite disease. Hum Mol Genet 19, R77–82

